# Biallelic loss-of-function *OBSCN* variants predispose individuals to severe, recurrent rhabdomyolysis

**DOI:** 10.1101/2021.06.04.447044

**Authors:** Macarena Cabrera-Serrano, Laure Caccavelli, Marco Savarese, Anna Vihola, Manu Jokela, Mridul Johari, Thierry Capiod, Marine Madrange, Enrico Bugiardini, Stefen Brady, Rosaline Quinlivan, Ashirwad Merve, Renata Scalco, David Hilton-Jones, Henry Houlden, Halil Aydin, Serdar Ceylaner, Jerry Vockley, Rhonda L Taylor, Hayley Goullee, Emil Ylikallio, Mari Auranen, Henna Tyynismaa, Bjarne Udd, Alistair RR Forrest, Mark R Davis, Drago Bratkovic, Nicholas Manton, Thomas Robertson, Pamela McCombe, Nigel G Laing, Liza Phillips, Pascale de Lonlay, Gianina Ravenscroft

## Abstract

Rhabdomyolysis is the acute breakdown of skeletal myofibres in response to an initiating factor, most commonly toxins and over exertion. A variety of genetic disorders predispose to rhabdomyolysis through different pathogenic mechanisms, particularly in patients with recurrent episodes. However, the majority of cases remain without a genetic diagnosis. Here we present six patients who presented with severe and recurrent rhabdomyolysis, usually with onset in the teenage years; other features included a history of myalgia and muscle cramps. We identified ten bi-allelic loss-of-function variants in the gene encoding obscurin (*OBSCN*) co-segregating with disease. We show reduced expression of *OBSCN* and loss of obscurin protein in patient muscle. Obscurin is proposed to be involved in SR function and Ca^2+^ handling. Patient cultured myoblasts appear more susceptible to starvation as evidenced by a greater decreased in SR Ca^2+^ content compared to control myoblasts. This likely reflects a lower efficiency when pumping Ca^2+^ back into the SR and/or a decrease in Ca^2+^ SR storage ability when metabolism is diminished. OSBCN variants have previously been associated with cardiomyopathies. None of the patients presented with a cardiomyopathy and cardiac examinations were normal in all cases in which cardiac function was assessed. There was also no history of cardiomyopathy in first degree relatives, in particular in any of the carrier parents. This cohort is relatively young, thus follow-up studies and the identification of additional cases with bi-allelic null *OBSCN* variants will further delineate *OBSCN*-related disease and the clinical course of disease.

## INTRODUCTION

Rhabdomyolysis is a serious medical condition involving the rapid breakdown of damaged or injured skeletal myofibres and may require intensive care management. Muscle breakdown results in release of myofibrillar content into the extracellular space and the circulation, resulting in hyperCKaemia (hyperCK) and myoglobinuria. Clinically, rhabdomyolysis can range from asymptomatic episodes with isolated hyperCK to a life-threating condition with profound myoglobinuria, often progressing to acute renal failure and requiring intensive care management in severe cases. Clinical features include acute muscle weakness, myalgia, muscle swelling and elevated CK (defined as >5-times the upper normal limit).^1^ Some patients also develop compartment syndrome necessitating fasciotomy.

Rhabdomyolysis can be acquired (trauma, ischaemia, infection and toxin or drug-related)^2^ or genetic^3^ in origin. A fascinating history of rhabdomyolysis exists in the literature, with reports dating back to biblical times and antiquity often in relation to poisoning.^4^ Quail poisoning is a well-documented cause of rhabdomyolysis and is caused by ingestion of toxins associated with quail meat.^4^

A schema has been suggested to discern rhabdomyolysis cases with a likely genetic contribution.^5^ This has an acronym of RHABDO (**R**ecurrent episodes, **H**yperCK for a prolonged period, **A**ccustomised physical activity, **B**lood CK >50xUNL, **D**rug/medication insufficient to explain severity, **O**ther family members affected/**O**ther symptoms.^5^ A large single-centre study showed that the most common triggers of rhabdomyolysis included ischaemia/anoxia and traumatic muscle injury in non-neuromuscular cases; whilst in >50% of cases with a known or suspected neuromuscular basis the trigger was exercise.^5^

There is a growing list of Mendelian gene defects associated with increased susceptibility to rhabdomyolysis, including a number of genes involved in muscle metabolism and mitochondrial function.^3; 6^ Recessive variants in the lipin-1 gene (*LPIN1*) are a common cause of childhood-onset and severe rhabdomyolysis, sometimes resulting in kidney failure and cardiac arrhythmia.^7; 8^ Impaired synthesis of triglycerides and membrane phospholipids have been hypothesised to underlie the pathogenesis of *LPIN1*-mediated rhabdomyolysis.

More recently, variants in genes encoding structural muscle proteins have been implicated in rhabdomyolysis. Bi-allelic variants in the gene encoding muscular LMNA-interacting protein (*MLIP*) have been shown to underlie a myopathy characterised by mild muscle weakness, myalgia, susceptibility to rhabdomyolysis and persistently elevated basal CK.^9^ Further, Alsaif *et al*. have reported a single case of rhabdomyolysis associated with a homozygous missense variant in *MYH1*, the gene encoding myosin heavy chain 2X^10^, and a *MYH1* missense variant is strongly associated with non-exertional rhabdomyolysis in Quarter horses.^11^

In addition, variants in known muscular dystrophy genes can also predispose a patient to rhabdomyolysis; in some cases rhabdomyolysis can be the presenting symptom of an underlying muscular dystrophy, e.g. *ANO5, CAV3 DMD, FKRP* and *SGCA*^5; 6; 12–15^ or neurogenerative disease, e.g. *TANGO2*.^5^

Variants in *RYR1* encoding the skeletal muscle Ca^2+^ release channel (Ryr1) of the sarcoplasmic reticulum (SR) have been increasingly recognised as an underlying cause of rhabdomyolysis.^16; 17^ Similarly, likely-pathogenic variants in genes critical to excitationcontraction coupling and Ca^2+^ handling (*CACNA1S* and *SCN4A*) have been implicated in exertional heat illness and rhabdomyolysis.^5; 18^ Kruijt *et al.* identified an underlying genetic diagnosis in 72 of 193 (37%) rhabdomyolysis probands that fulfilled one or more of the RHABDO criteria, these included variants in 22 disease genes.^5^ Despite sequencing of known rhabdomyolysis genes, including via large gene panels and exome sequencing, many individuals who experience rhabdomyolysis remain without a definitive genetic diagnosis.^5; 19–22^

Identification of the genetic cause in a patient with rhabdomyolysis is important because it enables appropriate advice on how to minimise future episodes, optimised clinical management and genetic counselling.

Obscurin is a component of the sarcomere and localises to the M-band and Z-disks.^23^ Obscurin interacts with titin, myomesin and small ankyrin 1 and is proposed to serve as a linker protein between the sarcomere and SR.^24–26^ Obscurin is also thought to be involved in SR function and Ca^2+^ regulation.^27; 28^ *Obscn* null (*Obscn*^-/-^) mice display a mild myopathy, including exercise-induced sarcomeric and sarcolemmal defects.^27–30^ Increased susceptibility of obscurin-deficient muscle to damage may trigger bouts of rhabdomyolysis in humans. Supporting this hypothesis, four of the six probands experienced rhabdomyolysis following exercise.

In this study we identified six patients with onset of severe recurrent rhabdomyolysis from 12-27 years of age and bi-allelic loss-of-function variants in the obscurin gene (*OBSCN*). Some patients had a history of myalgia and muscle cramps that preceded the initial episode of rhabdomyolysis. Between episodes, CK levels are normal to mildly-elevated. We showed reduced *OBSCN* transcript expression and protein abundance in muscle biopsies from affected individuals. Studies of patient cultured myoblasts showed that a starvation condition induces aberrant Ca^2+^ flux into the sarcoplasmic reticulum and higher levels of myoblast death under basal conditions, the hallmarks of the rhabdomyolysis.^31^

Our data clearly demonstrate that bi-allelic loss-of-function *OBSCN* variants predispose individuals to severe recurrent rhabdomyolysis. *OBSCN* variants should be considered in the diagnosis of patients with recurrent rhabdomyolysis.

## METHODS

All studies were approved by the Human Research Ethics Committee of the recruiting centre and all individuals participating in this study gave informed consent.

### Clinical Investigations

#### Patient details and investigations

We have clinically characterised six probands from six unrelated families originating from Australia (2), Finland, Turkey, the UK and the USA. Patients presented with severe rhabdomyolysis, from their teenage years. We performed pedigree analysis, neurological examination, including muscle strength evaluation according to the Medical Research Council (MRC) grading scale, serum CK levels during acute episodes and between episodes (baseline), lower limb muscle MRI and cardiac investigations in some cases, muscle biopsy and genetic workout in the probands and additional family members. The study was approved by the ethics committees of the participating institutions. Sample collection was performed after written informed consent from the patients according to the declaration of Helsinki.

#### Muscle pathology

Muscle biopsies were performed in all probands as part of routine diagnostic investigations. The samples were frozen in liquid nitrogen-chilled isopentane and processed for routine histological and histochemical techniques. Muscle samples were also collected for electron microscopy. Processing of muscle for light and electron microscopy was performed as outlined previously.^32^

### Genetic investigations

#### AUS1

DNA from the proband was run on version 1 of a custom designed neuromuscular disease targeted gene panel at Diagnostic Genomics, PathWest, as detailed in Beecroft *et al*.^22^ This did not identify any likely pathogenic variants. Whole exome sequencing was then performed using the Ion Proton^TM^ (Ampliseq chemistry, Life Technologies). Variant calling was performed using Torrent Suite V3.6.2. Data were annotated and filtered using an ANNOVAR annotation software suite. Pathogenicity predictions were made using online prediction software programs: SIFT, PolyPhen-2, and MutationTaster.

#### AUS2

DNA from the proband was sequenced on version 2 the PathWest neuromuscular disease gene targeted panel^22^. No likely causative variants were identified. DNA was subsequently re-sequenced on version 5 of the panel, which had been updated to include recently identified skeletal muscle disease genes, including *OBSCN*. All mapping and calling of variants was done by the BWA Enrichment App v2.1.2 on the Illumina Basespace Sequence Hub using our custom bed files. Data was analysed in Alissa Interpret (Agilent).

#### FIN1

Exome sequencing was performed on DNA from the proband as previously described^33^, and was analysed as a clinical exome that did not identify pathogenic variants in genes with previous disease associations in OMIM or ClinVar databases. This patient subsequently underwent targeted resequencing using the MYOcap gene panel.^34^

#### TUR1

Exome enrichment was performed using Twist Comprehensive Human Exome kit according to manufacturer’s instructions. Prepared library was sequenced on MGI DNBSEQ-G400 at 80-100X on-target depth with 150 bp paired-end sequencing at Intergen Genetic Diagnostic Centre (Ankara, Turkey). Bioinformatics analyses were performed using in-house developed workflow derived from GATK best practices at Intergen Genetic Diagnostic Centre.

#### UK1

Exome sequencing and analysis was performed on the proband’s DNA as outlined previously.^35^

#### USA1

Clinical exome sequencing was performed on the proband at GeneDX. No known or candidate pathogenic variants were identified. Re-examination of the exome in light of the association of *OBSCN* with recurrent rhabdomyolysis, identified a single heterozygous essential splice-site variant in *OBSCN*. DNA from USA1 was subsequently sequenced on version 3 of the PathWest neuromuscular disease gene panel. A second nonsense variant was identified in this individual, by the PathWest panel.

Bi-directional Sanger sequencing was used to confirm the *OBSCN* variants identified and where familial DNA samples were available, to show that the variants co-segregated with disease.

### Skeletal muscle RNA-seq data

Skeletal muscle RNA-seq data from a cohort of individuals were studied. These individuals included 30 patients with skeletal muscle disease, four patients with isolated hyperCK and one asymptomatic relative of a skeletal muscle disease proband. We utilised RNA-seq data that were generated using a ribodepletion method, this was to negate potential bias associated with RNA-sequencing of large genes, including obscurin, in samples generated with a poly-A RNA capture method.^36^

RNA was extracted with Qiagen RNeasy Plus Universal Mini Kit (Qiagen, Hilden, Germany) according to the manufacturer’s instructions. The strand specific RNAseq library was prepared using the Illumina Ribo-Zero Plus rRNA Depletion Kit (Illumina, Palo Alto, CA, USA) at the Oxford Genomics Center, Welcome Trust Institute, Oxford, United Kingdom. Sequencing was performed on Novaseq (Illumina), generating over 80 million 150bp-long reads per sample. Trimmed sequences were mapped against the hg19 human reference genome using STAR 2.7.0d.

To evaluate *OBSCN* exon usage, we analysed pooled junction data from the 35 RNA-seq experiments.

### Quantitative PCR

RNA was extracted from 30 mg frozen tissue using the RNeasy Fibrous Tissue Mini Kit (Qiagen) as described by the manufacturer. RNA was quantified with a Nanodrop ND 1000 spectrophotometer (Thermo Fisher Scientific) and electrophoresed on a 1% agarose gel to confirm RNA integrity and absence of genomic DNA contamination. The SuperScript III First-Strand Synthesis System (Thermo Fisher Scientific) was used to synthesize cDNA from up to 1 μg total RNA using random hexamers according to the manufacturer’s protocol. Prior to qPCR, all cDNA’s were diluted to the equivalent starting input of 100 ng RNA with UltraPure water (Thermo Fisher Scientific). The Rotor-Gene SYBR Green PCR Kit (Qiagen) was used to set up 10 μL reactions containing 1 μL diluted cDNA and 0.8 μM each of forward and reverse primers (*OBSCN, RYR1, ACTA1, MYOG, TBP, EEF2*; Supplementary Table 1). Primers were designed to amplify transcript variants 1 (NM_052843.4, isoform A), 2 (NM_001098623.2, isoform B), 3 (NM_001386125.1) and 1C (NM_001271223.2, inferred complete isoform). There were no predicted off-target products. There are two long noncoding RNA genes that overlap *OBSCN* and the primers do not amplify these. Primer efficiency (in the range 0.9 – 1.1) was validated by standard curve. Thermal cycling was performed on the Rotor-Gene Q real-time PCR cycler and data were analyzed with the associated software (Qiagen) using a cycle threshold of 0.03. Data were normalised to the geometric mean of two endogenous control genes (*TBP, EEF2*) using the delta-Ct method. Graphed data represent the mean ± SEM and were generated using GraphPad Prism (V6.02).

### Western blotting

Frozen muscle biopsies were homogenised in modified Laemmli sample buffer as described previously.^37^ Samples were run in Bio-Rad Criterion 3–8 % tris-acetate gradient gels (Bio-Rad Laboratories, CA, USA) at room temperature, 100 V, for 4 hours. The proteins were transferred from gels onto PVDF membranes with a Bio-Rad TransBlot Turbo device (program Standard SD, 60 min), using discontinuous buffer system, gel/anode buffer 1X CAPS with 0.1% SDS; PVDF/cathode buffer 1X CAPS. Subsequently, the post-blotting gels were stained with Coomassie Brilliant Blue, and the PVDF membranes were incubated in primary antibody solution, rabbit anti-obscurin ob59 (1:800 dilution) overnight at 8 °C (anti-obscurin domain 59 antibody is a gift from Prof. Mathias Gautel). Membranes were incubated with HRP-conjugated secondary antibody and the bands were detected using ECL (SuperSignal West Femto, Thermo Fisher Scientific) and ChemiDoc MP digital imager (Bio-Rad). Coomassie-stained gels were used to visualise titin and nebulin bands which served as size markers and loading controls.

### Cell-based studies

#### Immunofluorescence microscopy

Primary myoblasts were fixed and stained as previously described^38^ with anti-calnexin primary antibody (clone AF18, refsc-23954, Santa Cruz Biotechnology). Images were acquired on a confocal Leica LSM700 microscope, equipped with a 63X and a 1.3 numerical aperture (NA) oil immersion objective. Quantification and morphological analysis of endoplasmic reticulum was done with Icy v1.9.5.1 (BioImage Analysis Unit, Institut Pasteur, France).

#### Ca^2+^ measurements

Thapsigargin-induced responses were monitored in a FDSSμCell microplate reader (Hamamatsu Photonics, Japan) to assess sarcoplasmic reticulum (SR) Ca^2+^ content. Myoblasts were plated in 96-well plates at a density of 15,000 cells per well, and incubated for 1 hour in EBSS or in complete HAMF10 medium. Myoblasts were loaded with 4 μM Cal-520-AM (AAT Bioquest, CA, USA) for 45 minutes then washed in recording medium containing (mM) NaCl 116, KCl 5.6, MgCl_2_ 1.2, HEPES 20 (pH 7.3) and 150 μM EGTA. Thapsigargin (1 μM) was simultaneously added in the absence of external Ca^2+^ in all wells. Recordings were performed at 37 °C, frequency acquisition was 1 Hz and fluorescence signals (F) calibrated by adding 50 μM digitonin containing 6 mM Ca^2+^ (final concentrations) to obtain maximal fluorescence signals (F_max_). Data were expressed as F/F_max_ and SR Ca^2+^ content areas of the TG-evoked responses calculated using Origin software (OriginLab Corp, MA, USA). Thapsigargin was purchased from Alomone Labs (Israel) and all other reagents were from Sigma-Aldrich (France).

#### Apoptosis measurements

Myoblasts were plated in 96-well plates at a density of 5,000 myoblasts per well. Caspase 3/7 green apoptosis assay reagent and NuncLight red reagent for nuclear labelling (ref C10423 and 4717, Essen Biosciences Ltd, UK) were added in each individual well at a 1/1,000 final dilution. Four phase images and four fluorescent images per well (ex 440-480nm; em 504-544nm and ex 655nm; em 681nm) were taken using IncuCyte® S3 Life Cell Analysis System (Essen Biosciences Ltd, UK). Single green events representing caspase 3/7 positive myoblasts were counted and myoblast number was assessed from nuclear counts.

#### Statistical analysis

Statistical analysis was performed with GraphPad Prism software using a Mann-Whitney test for all image analysis experiments, Ca^2+^ and apoptosis measurements.

## RESULTS

### Clinical findings

Here we report six isolated patients presenting with a clinical picture of severe, recurrent rhabdomyolysis. Age-of-onset ranged from 12-27 years of age (median age: 17 years). In a consanguineous Turkish family, a younger brother (13 years of age) experiences myalgia and muscle cramps. There is no other family history of myalgia or rhabdomyolysis in any of the families reported in this study. Three of patients were/are elite-level athletes in their chosen fields: FIN1 was a competitive swimmer at a national level, USA1 is an elite high school lacrosse player and UK1 in early adult life competed nationally in the 200m and 400m distance races without symptoms. Detailed clinical summaries and investigations are available in a Supplementary document.

The triggers in two cases were exercise and heat (AUS1, TUR1), in two only exercise (AUS2, FIN1), in another both episodes occurred following travel (long-haul flight and bus trips; USA1) and in one the episodes occurred spontaneously (UK1). All patients experienced recurrent bouts of rhabdomyolysis (>two events), with one patient experiencing up to six episodes per year. Peak CK levels ranged from 17,000-603,000 IU/L (median 312,500 U/L). Basal CK ranged between normal to mildly elevated (<1,000 IU/L).

Other myopathic features are present between episodes of rhabdomyolysis, including myalgia (5/6), exercise intolerance (3/6) and muscle weakness (2/6). Three of the six probands experienced acute renal failure (AUS1, TUR1, USA1), in at least one case necessitating kidney dialysis. In two probands, rhabdomyolysis was associated with compartment syndrome (AUS1, USA1). In AUS1, compartment syndrome involved both lower limbs and required fasciotomy; in USA1 compartment syndrome during two episodes required fasciotomy.

None of the cases presented with cardiac involvement or have developed any symptoms of cardiac disease. There is no history of cardiomyopathy in first degree relatives of the probands.

AUS1 had resting tachycardia post presentation with rhabdomyolysis but no clear cardiac involvement. He had a normal cardiac MRI and normal echocardiogram. Cardiac investigations were also normal in UK1.

The clinical findings in each of the cases are presented in Table 1 and in the Supplementary clinical summaries. Lower limb muscle MRI in patients AUS1 and FIN1 (Supplementary Figure 1) were normal.

**Table 1:**
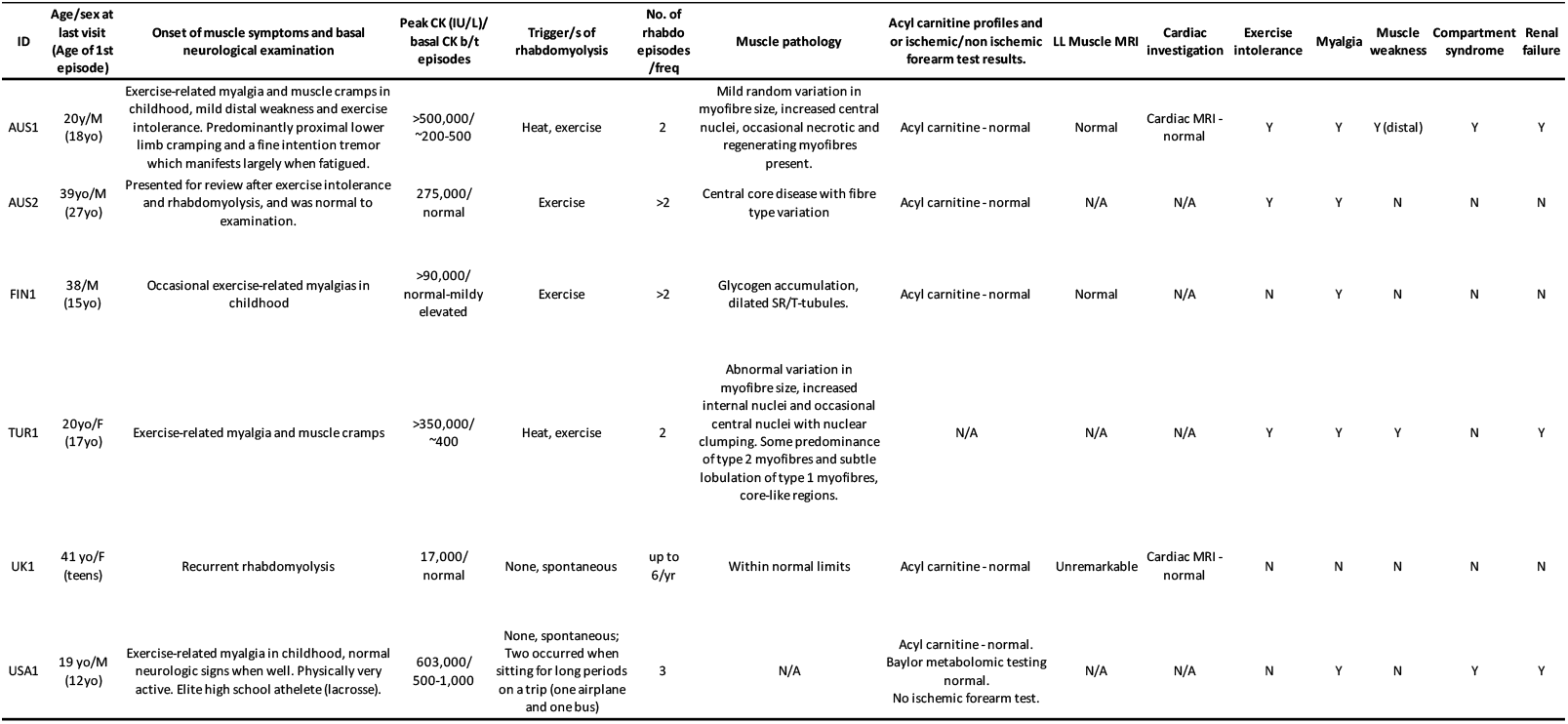
Clinical details of six probands with OBSCN variants.

Muscle biopsies were available for review in all cases. The findings on muscle biopsies ranged from within normal limits to non-specific mild myopathic changes and prominent central cores (Figure 2). Mild subsarcolemmal accumulations of glycogen (FIN1 and USA1), dilated SR and t-tubules (FIN1), mild increase in internal lipid droplets (TUR1), increased variation in myofibre size (AUS1, AUS2, TUR1, UK1), internal nuclei (AUS1, TUR1) and central cores in type I myofibres (AUS2).

**Figure 1:**
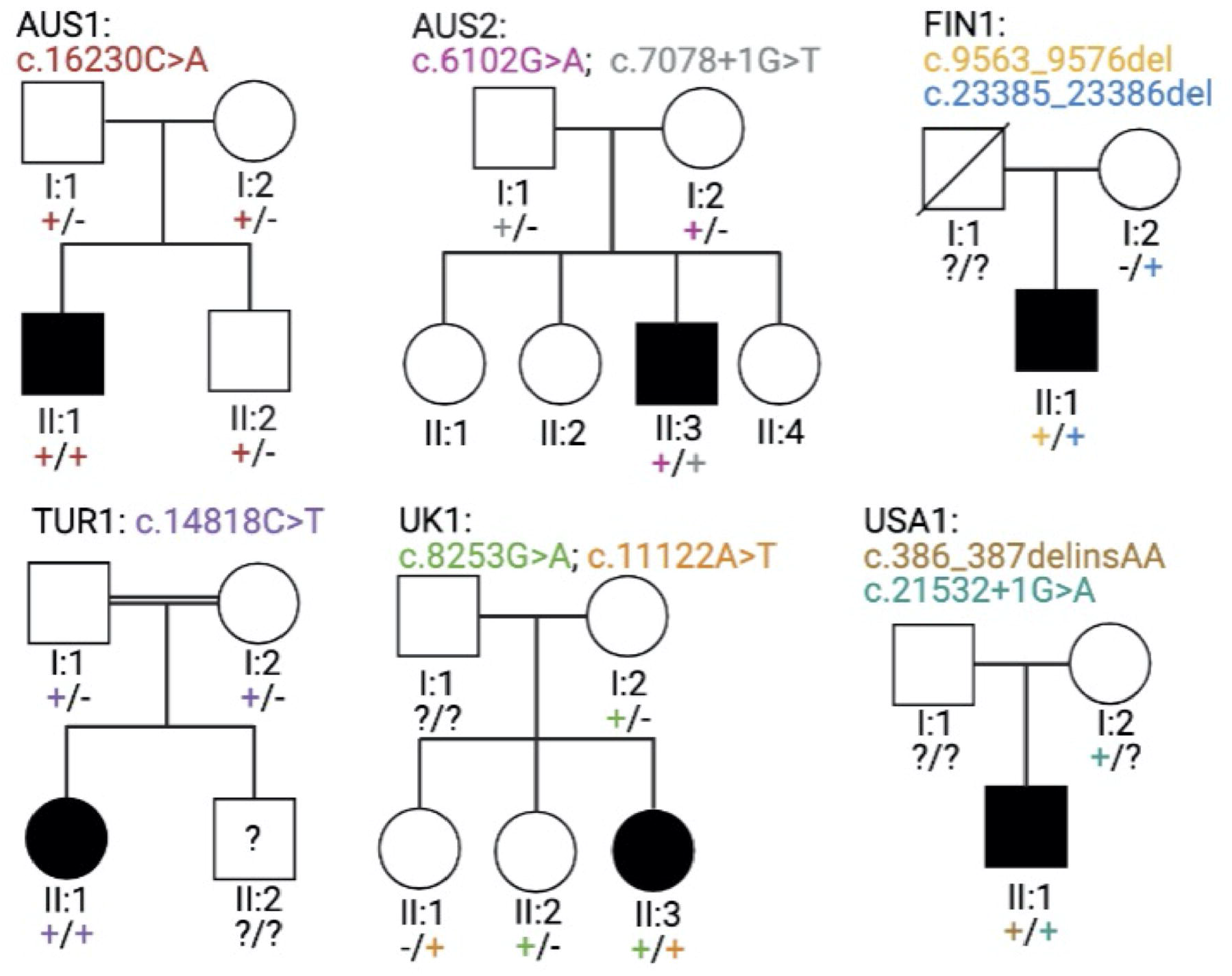
Pedigrees for the six families segregating bi-allelic loss-of-function variants in *OBSCN*.

**Figure 2:**
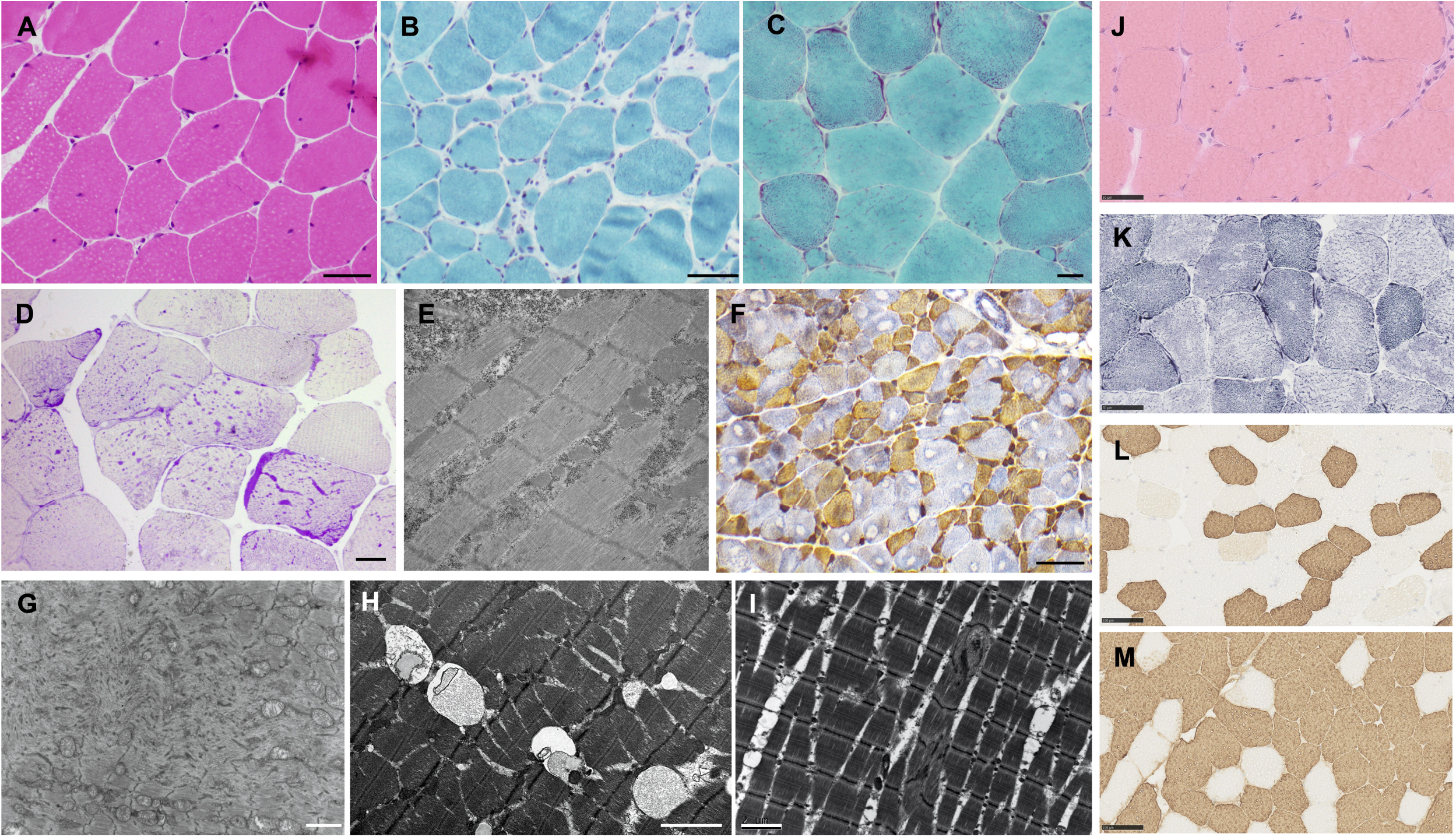
Muscle pathology, by light and electron microscopy (EM), associated with bi-allelic null *OBSCN* variants. Features ranged from within normal limits to mild myopathic changes with increased variation in myofibre size and internal nuclei (**A**: AUS1 and **J**: TUR1 [H&E], **B**: AUS2 and **C**: UK1 [Gomori trichrome]). Glycogen accumulations are evident in skeletal muscle from FIN1 (**D**; periodic acid-Schiff staining) and USA1 (**E**, EM). Prominent central cores are seen in type I myofibres of AUS2 (**F**, NADH(blue)-fast myosin(brown) combined enzyme-immunohistochemistry and **G**, EM). **K**: NADH staining on TUR1 shows mild disruption of internal architecture, type I myofibres appear almost lobulated and occasional small core-like areas. Slow (**L**) and fast (**M**) myosin staining in TUR1 shows mild predominance of type II myofibres, type I myofibres relatively small and few intermediate myofibres.Dilated SR and t-tubules are also evident in the muscle biopsy of FIN1 (**H**; EM). Focal Z-band streaming is seen in skeletal muscle from AUS1 (**I**; EM). Scale bars: 20 μm (C, D), 50 μm (A, B, K, J) 100 μm (F, L, M), 2 μm (G, H, I).

### Genetics

We identified ten rare or novel bi-allelic loss-of-function variants in *OBSCN* co-segregating with rhabdomyolysis in six families (Table 2; Figure 1). In *AUS1* we identified a homozygous nonsense variant in *OBSCN* (NM_001271223.2 corresponds to the inferred complete (IC) obscurin isoform; exon 62, c.16230C>A, p.(Cys5410*)). The variant (*rs*1322344930) is present on four of 273,940 alleles in gnomAD. The variant was confirmed by bi-directional Sanger sequencing; studies of familial DNA showed that both healthy parents and younger brother were all carriers of the variant. In AUS2 we identified two variants, a nonsense variant in exon 21 (c.6102G>A, p.(Trp2034*)) and an essential splice donor site variant (exon 24, c.7078+1G>T). The nonsense and essential splice-site variants were maternally and paternally inherited, respectively. This essential splice-splice variant is predicted to remove exon 24 resulting in an in-frame deletion of 89 amino acids [p.(Val2270_Arg2359del)]. We did not have access to RNA-seq data from AUS2, however, we have data from an unrelated patient that also carries this variant. RNA-seq studies on skeletal muscle found that the main consequence of the c.7078+1G>T variant is skipping of the first two nucleotides of exon 25 (data not shown). Thus, the major consequence of this change is likely a frameshift. Both were rare in gnomAD (allele frequency <0.0002), with no homozygotes present. Patient FIN1 harboured bi-allelic *OBSCN* deletions; exon36: c.9563_9576del and exon105: c.23385_23386del, p.(Ser7796*). The c.9563_9576del variant is novel, whilst the c.23385_23386del is present on 740 alleles in gnomAD including four homozygote individuals. Three of the four homozygotes are of Finnish background and the allele frequency in Finns is 0.006. In TUR1 a homozygous rare nonsense variant was identified in exon 46 (c.14818C>T, p.(Arg4940*)). This variant was present on five of 209,424 alleles in gnomAD, there were no homozygotes. Sanger sequencing found that each parent carried the variant. The variant was absent from ~2,500 Turkish exomes suggesting that this is not a common variant in the Turkish population. UK1 harboured bi-allelic nonsense variants (exon 31: c.8253G>A, p.(Trp2751*) and exon 42: c.11122A>T, p.(Lys3708*)). Two healthy siblings had single mono-allelic variants. The c.8253G>A variant is novel and the c.11122A>T variant is present on two alleles in gnomAD. USA1 harboured a multinucleotide polymorphism (MNP) in exon 2 which is annotated as c.386T>A, p.Phe129Tyr (*rs*749567826) and c.387C>A (*rs*769050588), p.Phe129Leu but is c.386_387delinsAA, p.(Phe129*). This MNP is present on 114 alleles in gnomAD, including two homozygotes. Both homozygotes are within the Ashkenazi Jewish population and the allele frequency in this population is 0.012. The second variant in USA1 occurs at the essential splice donor site of exon 90, c.21532+1G>A; this rare variant is present on three alleles in gnomAD. The exon 90 splice-site variant is inherited maternally and is predicted to result in skipping of exon 90 and a frameshift deletion.

**Table 2:**
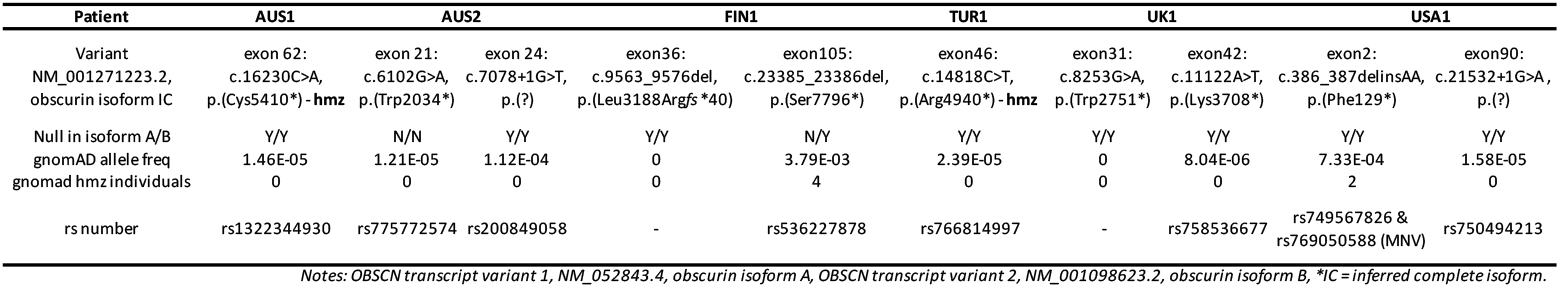
Details of the OBSCN variants identified in six recurrent rhabdomyolysis probands.

### Obscurin exon usage

In skeletal muscle, *OBSCN* encodes two canonical isoforms (A and B); obscurin A (~720kD) includes 65 immunoglobin domains and two fibronectin III domains along with a number of C-terminal signalling domains.^39^ The larger isoform, obscurin B (~870kD), has a similar structure to obscurin A but diverges at the C-terminal region where it contains two Ser/Thr kinase domains^39^. Exon 21 and 105 are annotated in the IC obscurin isoform, however exon 21 is not present in obscurin A or B and exon 105 is only present in the short isoform (isoform A). To investigate this, we examined exon usage in RNA-seq data from human adult skeletal muscle samples. Exon 21 was present in 93% (80,165 reads supporting the junction between exons 20 and 21) of all transcripts (Figure3A). The shorter isoform of obscurin (obscurin A) utilises an alternative 3’UTR in exon 92. RNA-seq analysis showed that 61%of all transcripts correspond to this short isoform (90070 reads supporting the inclusion of the exon 92 alternative 3’UTR). There were 39% of reads (*n*=58,551) supporting the skipping of the alternative 3’UTR in exon 92, and accounting for the longer isoform (obscurin B; Figure 3A).

**Figure 3:**
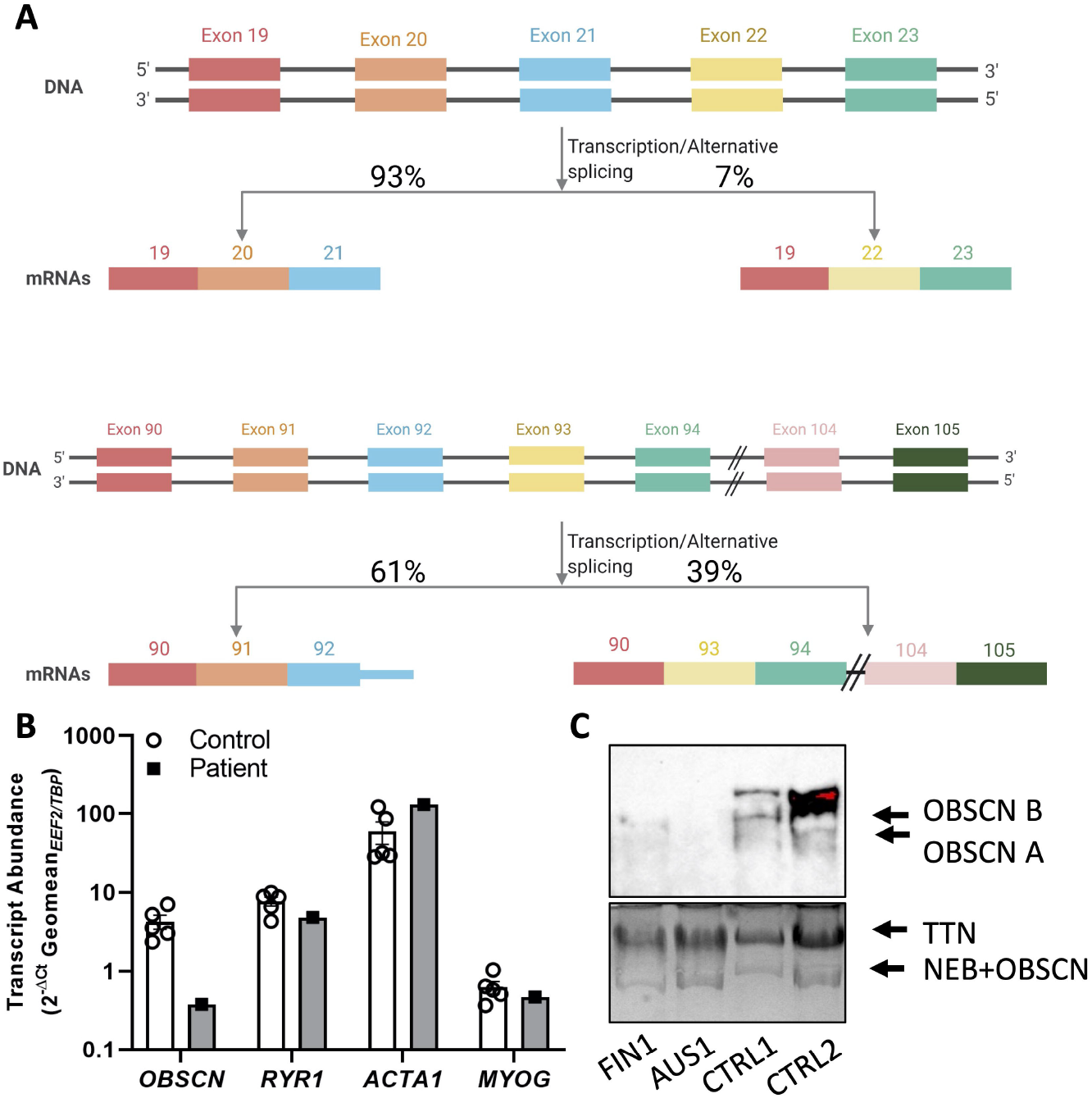
Obscurin transcript and protein abundance in healthy control and patient skeletal muscle. (**A**) A schematic showing exon usage of exon 21 and 105, generated from skeletal muscle RNA-seq data. (**B**) Transcript abundance of *OBSCN, RYR1, ACTA1* and *MYOG* in human skeletal muscle obtained from the patient (filled square, grey bars) and five unrelated controls (open circles, white bars) was assessed by qPCR. *OBSCN* transcript abundance is specifically reduced by more than 10-fold in AUS1 patient skeletal muscle relative to controls. Expression of each transcript was normalised to the geometric mean of two endogenous control genes (*EEF2* and *TBP*) using the delta Ct method. Graphed data represent the *mean ± SEM*. (**C**) Western blot showing reduction/absence of OBSCN in patient muscle (FIN1, AUS1) compared to healthy control (CTRL) samples. Coomassie staining of the TTN and NEB protein bands are shown to demonstrate loading of total muscle protein.

Similarly, at a protein level, the ratio between obscurin A and B isoforms is almost 50-50 in the adult muscles (Figure 3C). We therefore conclude, based on knowledge of *OBSCN* expression, that each of the variants identified is predicted to result in loss of a considerable portion of obscurin A and/or B isoforms.

### Obscurin transcript and protein levels are reduced in patient skeletal muscle

To determine whether the homozygous AUS1 variant (c.16230C>A, p.(Cys5410*)) was associated with a decrease in *OBSCN* transcript abundance, we performed qPCR using cDNA obtained from patient muscle biopsy and five unrelated control muscle biopsies. The average normalised *OBSCN* transcript abundance of control samples was 11.2-fold greater than in the patient muscle (average normalised OBSCN transcript abundance of 4.25 ± 1.9 in the control, versus 0.38 in the patient; Figure 3B). To ensure that this difference was specific to *OBSCN* transcript, and not an artifact, we also measured the transcript abundance of three additional genes; ryanodine receptor 1 (*RYR1*), skeletal muscle alpha-actin (*ACTA1*) and myogenin (*MYOG*). For each of these three genes, the transcript abundance measured in the patient sample was within the range of values obtained for the control samples (Figure 3B). This indicates that the decreased *OBSCN* transcript abundance is likely to be a real finding.

Western blot performed for obscurin in skeletal muscle from FIN1 and AUS1 showed greatly reduced levels of both isoforms of obscurin (A and B) compared to three healthy control muscle samples (Figure 3C). In FIN1 there is some retention of obscurin A, this most probably represents obscurin A arising from the allele harbouring the nonsense variant in exon 105 that is excluded in the short isoform A. This suggests that the disease manifests due to reduced levels or absence of obscurin protein in patient skeletal muscle. Total loading of muscle protein is indicated by band intensities for other large muscle proteins (titin and nebulin).

### Ca^2+^ handling is impaired in cultured patient cells

Ca^2+^ is tightly regulated in skeletal muscle and impairment in Ca^2+^ channel function is a recognised mechanism in rhabdomyolysis. Moreover, obscurin is also thought to be involved in SR function and Ca^2+^ regulation.^27; 28^ To analyse the SR network, we performed immunolabeling with the antibody anti-calnexin on primary myoblasts from a healthy control and patient UK1, in growth medium. By confocal microscopy, we did not find any differences in patient myoblasts compared to the control (Figure 4A), not even after quantification of myoblast total SR nor in other morphological parameters (sphericity, length (Figure 4B), roundness, elongation and flatness (not shown)).

**Figure 4:**
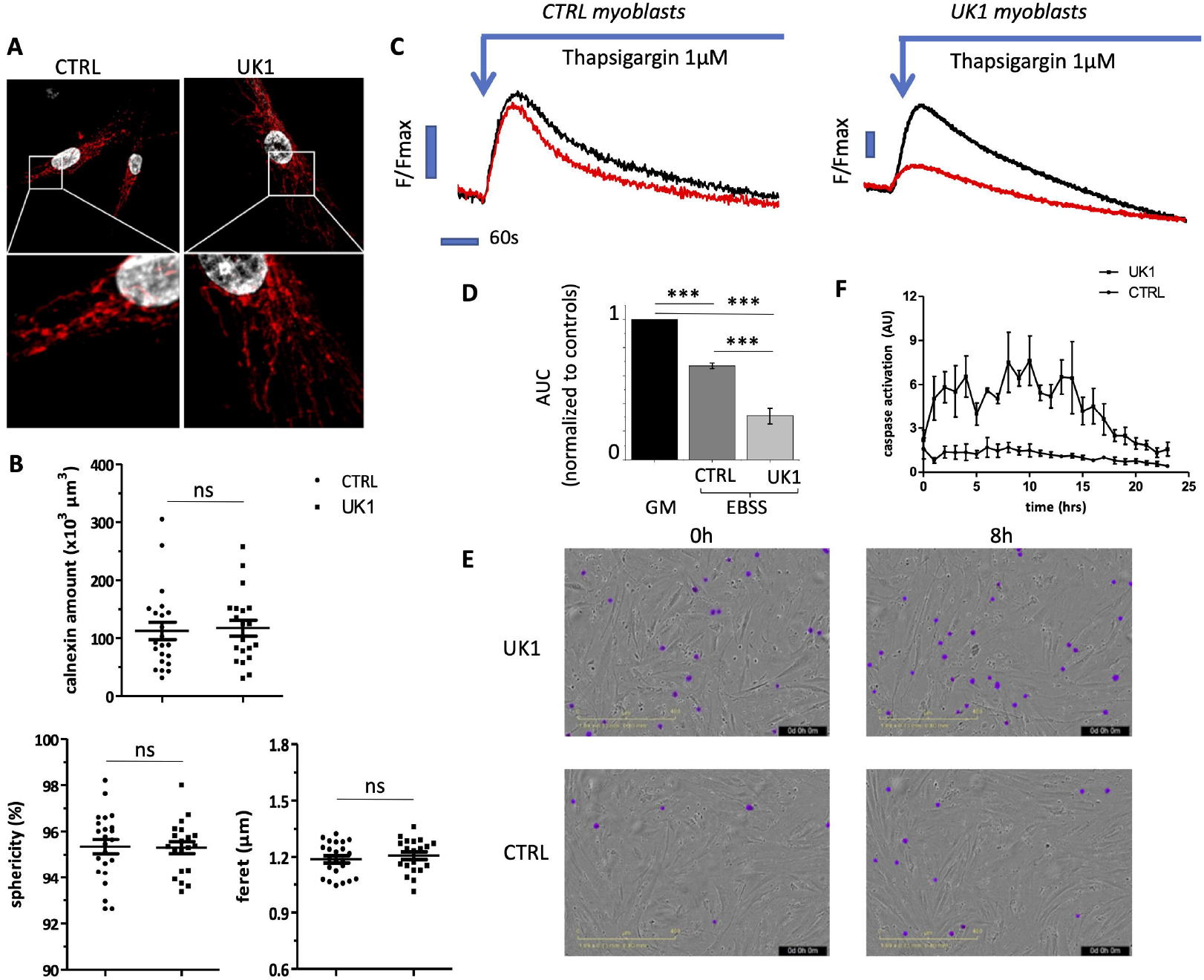
Studies from patient (UK1) myoblasts show aberrant Ca^2+^ flux and increased cell death. **(A)** SR morphology is not altered in patient myoblasts when compared to control myoblasts as shown by immunostaining with anti-calnexin antibody and confocal analysis. (**B**) Total SR content and the morphological parameters measured (length, sphericity) are similar in healthy control (CTRL) and patient myoblasts. **(C)** Representative SR Ca^2+^ content measurements in myoblasts from a healthy control (CTRL, left panel) and patient (right panel) in control (GM, black traces) and EBSS medium for 2 hours (red traces). (**D**) SR Ca^2+^ content was assessed from the area under the curves (AUC) after thapsigargin addition. Horizontal bar, 0 seconds; vertical bars F/Fmax 0.1 (arbitrary units). Histograms summarising area under the curve (AUC) in control (GM) and EBSS media. Data from 12 and six individual wells for CTRL and patient respectively obtained from two independent experiments corresponding to a decrease of 33±2 % (CTRL) and 69±6 % (Patient) of SR Ca^2+^ contents in EBSS medium. (**E**) Apoptosis was assessed from purple events representing caspase 3/7 positive cells normalised to cell number. Patient myoblasts show higher levels of apoptosis (2.3-fold) as detected by caspase 3/7 expression when compare to CRTL myoblasts. (**F**) Quantification of caspase 3/7 expression in control and patient myoblasts. Results of one representative experiment out of two independent experiments.

We studied regulation of SR Ca^2+^ content in myoblasts and found that starvation (EBSS medium) induced a decrease in Ca^2+^ SR content when compare to normal growth conditions (Figure 4C, p<0.001) probably reflecting lower efficiency when pumping Ca^2+^ back into the SR or a decrease in Ca^2+^ SR storage ability when cell metabolism is diminished. Susceptibility to starvation is exacerbated in UK1 myoblasts as we observed a 69±6%decrease in Ca^2+^ SR content compared to 33±2% in control myoblasts (Figure 4D, p<0.001). This result suggests that patient myoblasts have a decreased ability to fill the SR during starvation conditions for the same reasons as described above. Moreover, obscurin deficiency is associated with greater myoblast death under basal conditions as attested by caspase expression (Figure 4E, p<0.001). When quantified by flow cytometry, percentage of cell death in patient myoblasts was 54% compared to 34% in control myoblasts (Figure 4F).

## DISCUSSION

Herein we described six patients with susceptibility to severe, recurrent rhabdomyolysis (peak CKs ranged from 17,000-603,000 IU/L) due to bi-allelic loss-of-function variants in *OBSCN*. All cases had experienced at least two episodes of rhabdomyolysis, with one individual (UK1) experiencing up to 6 episodes per year. Triggers included exercise (including mild exercise, n=4) and heat (n=2); in two individuals the episodes occur without obvious triggers.

In most cases there was a prior history of myalgia and muscle cramps. Basal CK between episodes ranged from normal to mildly elevated (<1,000 IU/L). Three patients were/are elite level athletes in their chosen fields. Elite athletic performance preceding the onset of neuromuscular disease has been noted anecdotally for other genetic neuromuscular diseases; most notably in patients with pathogenic variants in *ANO5, CAPN3, CAV3, DYSF* and *RYR1*.^3; 14; 40–43^

Features on muscle biopsy ranged from unremarkable and minimal non-specific changes through to striking central cores in type I myofibres. Increased internal nuclei, mild variation in myofibre size and glycogen accumulations were each reported in more than one case. Other features included increased lipid droplets, core-like regions, type II myofibre predominance and occasional myofibre atrophy and nuclear clumps.

Population variant frequency data shows that all LOF variants for which there are homozygous individuals in gnomAD are flagged as a MNV (which together do not result in LOF changes at the amino acid level) or have LOF curation notes of “*uncertain*” or “*not LOF*” (Supplementary Figure 2). This suggests that bi-allelic LOF variants in *OBSCN* are likely to be pathogenic. This association was perhaps delayed due to the presence of a relatively common spurious LOF variant in *OBSCN* (p.Arg3252* [AGA>TGA], *rs*3795786, allele frequency: 0.03, gnomAD: >900 homozygote individuals). However, this variant was subsequently annotated as a MNP associated with a protein change of p.Arg3252Leu (AGA>TTA). One of the variants (p.(Ser7796*)) we identified in exon 105 is present in four individuals in the homozygous state in gnomAD and is present on one allele in FIN1. This annotation of “*uncertain*” is likely due to this exon not being predicted to cause a null allele in OBSCN isoform A. However, we have been able to show by RNA-seq that this exon is expressed at similar levels to flanking exons present in both major isoforms of OBSCN. Furthermore, by western blot we have shown loss of OBSCN isoform A and B in FIN1 skeletal muscle.

Two variants reported in this cohort are present in ~1% of individuals in specific control populations and thus these likely founder variants may underlie other cases of rhabdomyolysis in these populations. The MNV (c.386_387delinsAA, p.(Phe129*)) identified in USA1 is present at an allele frequency of 0.012 in the Ashkenazi Jewish population and there are two homozygous Ashkenazi Jewish individuals in gnomAD. The p.(Ser7796*) variant present in FIN1 is present at an allele frequency of 0.009 in the Finnish population in gnomAD.

The finding of similar expression of exons not thought to be included in some isoforms of *OBSCN* is similar to the identification of exons within *TTN* that were thought to only occur in the meta-transcript but were later shown to be present in *TTN* transcripts in adult skeletal muscle.^44^ Our findings suggest that further studies are needed to provide a comprehensive picture of the complex *OBSCN* splicing pattern. As already demonstrated with the even larger *TTN* transcripts, this is crucial for a proper clinical interpretation of variants in such large genes.^45^

Obscurin was originally identified as a titin-binding protein and has been observed to localise to the M-band and also the Z-disk of striated muscle.^25^ At the M-band, obscurin interactions with titin, myomesin and myosin binding protein C. Binding at the titin C-termins Ig domain (M10) is responsible for the predominant M-band localisation in mature myofilaments. Heterozygous variants in the titin M10 domain cause dominant tibial muscular dystrophy (TMD)^46^ and bi-allelic variants cause LGMD R10.^47^ These variants disrupt binding with obscurin. TMD variants associated with different clinical severity, correlate with the degree of loss of obscurin interaction.^25^

Four obscurin isoforms have been characterised, including two high-molecular weight proteins (obscurin A and B) that are abundant in skeletal muscle.^28^ Obscurin A contains two COOH-terminal binding sites that can interact with ankyrin proteins^48^, including small ankyrin 1 (sAnk1.5) of the sarcoplasmic reticulum (SR). Obscurin is thus proposed to play a key role linking the contractile apparatus to the SR. *Obscn* null^27^ and *sAnk1.5* null^49^ mice both show reduced longitudinal SR volume. Muscle from patient FIN1 showed markedly dilated T-tubules by electron microscopy, this may indicate that the SR is impaired in obscurin-related myopathy.

Obscurin’s precise role in skeletal muscle development, function and disease has remained *obscure*.^50^ It had been shown in *C. elegans, D. Melanogaster* and *D. Rerio* that obscurin and its homologue unc-89 might play an important role in sarcomerogenesis and the lateral alignment of sarcomeres.^51–53^

However, studies by Lange and colleagues of *Obscn* null mice failed to identify any defects in sarcomere structure and alignment.^27; 28^ In contrast, Sorrentino’s team also generated and studied *Obscn* null mice, and found defects in sarcolemmal integrity and muscle damage in response to exercise.^29; 30^ No defects suggestive of rhabdomyolysis or impaired Ca^2+^ handling were noted in these mice studies; however it is tempting to postulate that muscle damage triggered by obscurin deficiency, may lead to rhabdomyolysis susceptibility. Indeed, four of our six patients experienced rhabdomyolysis following exercise, in some instances mild exercise was sufficient to trigger an episode, i.e. climbing a flight of stairs.

Our patients are relatively young, it will be interesting to follow up this cohort and identify additional cases with obscurin deficiency to determine the natural history and progression of the disease into later life.

Heterozygous variants (missense, splice-site and frameshift) in *OBSCN* have been associated with cardiomyopathy (dilated cardiomyopathy, hypertrophic cardiomyopathy, left ventricular non-compaction) in patients.^39; 54; 55^ However, as highlighted in Grogan and Kontrogianni-Konstantopoulos the functional consequences of the identified *OBSCN* variants remain elusive.^56^ *Obscn*^-/-^ mice do not exhibit any signs of cardiomyopathy.^27^ In GTEx, *OBSCN* is highly enriched in skeletal muscle (median TPM 213) compared to all other tissues and is expressed at much lower levels in the heart (left ventricle: median TPM 35, atria appendage: median TPM 24; https://www.gtexportal.org/home/gene/OBSCN). In a systematic review of cardiomyopathy genetics, Ingles *et al.* classified the association between *OBSCN* variants and dilated cardiomyopathy as “limited” based on the available literature and scientific evidence.^57^ None of the patients in this cohort nor any of their carrier first-degree relatives report any cardiac involvement.

In summary, we have identified bi-allelic loss-of-function variants in *OBSCN* as a cause of recurrent rhabdomyolysis, typically presenting in teenage years. *OBSCN* should be considered in the genetic diagnosis of rhabdomyolysis.

## ACKNOWLEDGEMENTS

This work is supported by NHMRC grants (APP1080587, APP1146321 and APP2002640) to GR, ARRF and NGL. GR is supported by an NHMRC CDF (APP1122952) and NGL and ARRF are supported by NHMRC Senior Research Fellowships (APP1117510, APP1154524). This work was also supported by the Academy of Finland Neurogenomics pHealth funding. The authors thank Nicolas Goudin and Meriem Garfa from the cell-imaging platform; the Oxford Genomics Centre at the Wellcome Centre for Human Genetics (funded by Welcome Trust grant reference 203141/Z/16/Z) for the generation and initial processing of the RNA sequencing data. This work was supported by grants to PdL from Fondation maladies rares, Agence Nationale de la Recherche (ANR – AAPG 2018 CE17 MetabInf), the Association Française contre les Myopathies (AFM 2016 – 2018 19773), and patient associations (Nos Anges, AMMI, OPPH, TANGO2 family associations, Hyperinsulinisme).

