## Supplementary Information for "Biallelic loss-of-function *OBSCN* variants predispose individuals to severe, recurrent rhabdomyolysis"

**Supplementary clinical details**

***AUS1***: He first came to medical attention at the of age 11 years with lower limb muscle cramping and CK ~ 4,000 IU/L following a week of school swimming lessons. At that time recurrent lower limb cramping from early childhood was reported. There was mild distal weakness, exercise intolerance and fine intention tremor. From that time, the patient had recurrent episodes of acute increases in CK following exercise. His first hospitalisation for rhabdomyolysis was at the age of 18 years after swimming in warm surf in Northern Australia. Subsequently he presented in 2017 with profound rhabdomyolysis (CK> 500,000 IU/L) at the age of 20 years following exposure to heat while walking around his university campus in South Australia. This presentation was complicated by compartment syndrome requiring bilateral fasciotomy in his proximal and distal lower limbs, and acute kidney injury requiring dialysis. He had a prolonged ICU stay and required rehabilitation but was able to walk unaided on discharge. He has some residual mild peripheral neuropathy. A resting tachycardia has been noted since his admission, however recent cardiac MRI was normal as was lower limb MRI of his musculature. Prior to his presentation in 2017 his baseline CK was ~ 200 IU/L but this increased to ~ 500 IU/L. Chronic symptoms in AUS1, include predominantly proximal lower limb cramping and a fine intention tremor which manifests largely when fatigued.

Muscle biopsy (right quadriceps at 15 years of age) revealed increased central nuclei and fibre size variation. Occasional necrotic and degenerating myofibres were seen. On electron microscopy, features included focal Z-band streaming and a subsarcolemmal filamentous body.

***AUS2***: Currently aged 40 years. AUS2 presented to medical care in 2008 with severe rhabdomyolysis (CK= 275,000 IU/L) after an intensive spin class course over several days. He developed widespread muscle pain, then noted dark urine and sought medical attention. There had been other episodes of rhabdomyolysis with lesser elevations of CK.

He had been very fit and active and played Australian football and soccer and did weightlifting. He had avoided long distance running because of fatigue and lack of endurance. As a child he had been thought to have hypotonia and muscle pain. He also had episodes of loss of muscle tone, with falling to the ground. His parents and three sisters were unaffected.

He had two anaesthetics without complications.

He does not take any medication, but used creatine and whey protein supplements.

On examination there was good muscle bulk, and normal power, tone and reflexes.

Since the episode of rhabdomyolysis, he has modified his exercise regimen and there have been no further episodes.

A muscle biopsy was taken when he was well and some months after an episode of rhabdomyolysis. The muscle biopsy showed central cores in type I myofibres. There was inflammation to a degree that could not be dismissed.
Initial genetic screening found no causative mutation in *RYR1*.

***FIN1***: The patient was first hospitalised because of rhabdomyolysis (CK 69,000 IU/L) provoked by a swimming workout at age 15. He had occasional exercise-related myalgias in childhood, but was otherwise healthy and could swim and run on a competitive level. A year after the index episode, he had another exercise-related rhabdomyolysis (CK >90,000 IU/L). Over subsequent years, he has had several CK elevations of >7,800 IU/L (the highest measurable level in a local laboratory) combined with lower limb myalgias provoked by seemingly innocuous triggers, such as running down one flight of stairs. CK values remained normal or only mildly elevated, when the patient avoided exercising apart from normal daily walks at an average pace. Over the years, lower limb muscle MRI (Figure Supplementary 1) and EMG/NCS have been performed several times with normal results. A bicycle spiroergometry exercise test showed normal lactate elevation at age 34 years and blood acylcarnitine levels were within reference range. More recently, the patient has been able to continue regular endurance training 5 days a week without further muscle symptoms over the last two years. It is unclear whether this improvement might be age-related or due to changes in diet or exercise regimen the patient has reportedly implemented, or some other unknown factors.
Two previous muscle biopsies were reportedly unremarkable. A third muscle biopsy, obtained from *vastus lateralis* muscle when the patient was 38 years old, displayed mild intermyofibrillar and subsarcolemmal accumulation of glycogen and dilated sarcoplasmic reticulum/T-tubule structures (Figure 2).

***TUR1***: The first symptoms of the patient were long-standing muscle cramps, leg pains, and exertional myalgia after walking short distances starting around the age of six years. These symptoms were enough for her to take days off school. She was first hospitalised for rhabdomyolysis (CK >350,000 IU/L) with acute renal failure, provoked by playing basketball at age of fourteen year and six months. She stopped playing basketball and other games requiring physical activity after the first episode because of pain attacks in her legs, gluteal and lumbosacral areas lasting one or two days. She also complained of stiffness around muscles of the lower extremities in addition to pain after physical activities. At the age of sixteen and 6 months, she had one episode of vomiting illness for which she attended a local emergency department and her CK level was found to be 400 IU/L, she recovered from this episode quickly without any obvious complications after increasing her oral intake appropriately. The second episode of severe rhabdomyolysis, requiring three-week hospitalisation, occurred at the age of seventeen and one months after long-distance walking. One the first day of that episode, she described cramps and pain in the thighs, lumbosacral and gluteal region. The pain got worse not allowing her to move, and she felt dizziness on the second day of the episode. She was admitted to the hospital after her urine turned red-brown colour on the third day. Her CK level was found to be 33,100 IU/L at admission, her urine output was decreased but there was not any biochemical evidence of renal insufficiency. Fluid replacement therapy and paracetamol, morphine sulfate and diazepam for pain were given. The CK level increased to 140,327 IU/L on the second day of the hospitalisation and decreased to 1,894 IU/L by the 7th day of the hospitalisation. The urine discolouration of myoglobinuria lasted about two days after fluid replacement, and the pain took about one and a half weeks to resolve.

Muscle biopsy performed after the second episode showed abnormal myopathic features including abnormal variation in myofibre size with some small fibres and a little hypertrophy (up to 100 microns) with increased in internal nuclei and occasional central nuclei with nuclear clumping, and some predominance of type 2 myofibres with lobulation of type 1 myofibres. lmmunohistochemistry for dystrophin, sarcoglycan, laminin alpha2 and desmin were found to be normal. Blood acylcarnitine profile and fatty acid oxidation studies on fibroblasts were normal. Muscle respiratory chain enzymes complex 1 to 4 were normal.

Dystrophin MLPA and full gene sequencing of *DMD*, *FKRP, ANO5*, and *DYSF* did not identify any likely pathogenic variants. A heterozygous *CACNA1*S variant (c.4546G>T p.(Asp1516Tyr) was identified in the patient and her unaffected mother.

She had an uncomplicated abdominal liposuction surgery, under general anaesthesia, at the age of 19 years. She has not had any severe rhabdomyolysis event for the last three years, but she has pain and cramps, even after 10 minutes of walking.

***UK1***: The UK1 patient, had at least two muscle biopsies. One performed in 1998 at Queen Square which was unremarkable but for the presence of rimmed vacuoles (documented in a letter). A second biopsy was performed in Oxford in 2014 and this showed myopathic changes (see Figure 2).
Symptoms started in adolescence. Since then has had up to 6 attacks a year, lasting between 2 days and 3 weeks. There is no obvious precipitant to the attacks. Specifically, there is no certain relationship to exercise. In her early adult life she was a high-performing athlete – running 200 and 400 metres at a national level. Such activity never precipitated an attack. She now exercises regularly, but at a lower level of intensity. She had wondered if an attack would sometimes be precipitated by

a slight change in her routine. There is no clear relationship between the attacks and food intake. She has always eaten a good diet in a highly regular fashion. If she goes on holiday that pattern may change slightly, and she had wondered if that might be a factor. There is no history to suggest that any of her attacks have been precipitated by fasting.
Attacks present as rapid onset of widespread muscle pain. She has never noticed myoglobinuria. CK recorded at up to 17,000 IU/l in attacks. Normal CK between attacks.
No other medical history of note. No family history of note.
Examination between attacks normal.
Normal exams: Acylcarnitines, muscle biopsy (report).

***USA1***: This is a now 19-year-old male with a history of recurrent rhabdomyolysis. He was well until age 12 years when he had an initial episode of muscle pain in the lower extremity following a long plane trip. A year later he had similar pain during sports practice. This resolved after one week of rest with no medical intervention. He experienced his first episode of acute rhabdomyolysis at 12 years of age, following the start of cross-country season at school and a long car trip. His maximum CK was 397,580. During that hospitalization he had a fasciotomy on both thighs. He experienced ventricular tachycardia with elevated potassium levels but did not need dialysis. He had two additional severe episodes at age 14 years. During one, his maximum CK was 603,000. He had another fasciotomy performed on the right leg and received plasmapheresis for kidney failure. His maximum creatinine was 4.3 and his maximum BUN was 59. He was subsequently treated with Dantrolene and has had no further episodes of acute rhabdomyolysis, though his resting CK levels are often in the 500-1,000 range. His physical exam when well is unremarkable, with normal muscle bulk, tone and strength. He does not have other neurologic signs. Metabolic testing was not performed when he was acutely ill, but urine organic acids, and blood lactate, ammonia, acylcarnitine profile, and amino acids have been normal at multiple times when well. Undirected metabolomic testing through Baylor School of Medicine Laboratory failed to identify any significant abnormalities.

**Supplementary Table 1. Primers used in qPCR.**

| Gene (transcript) | Forward (5' – 3') | Reverse (5' – 3') |
|---|---|---|
| *OBSCN* (NM_052843.4) | CTGCTAGTGCTGGTGATCCG | GCAGCATGGGGCTCTGAATG |
| *RYR1* (NM_000540.2) | TGGTGGACATGCTCGTGGAA | CGCTGAACTGCTTCTGGCTG |
| *ACTA1* (NM_001100.3) | AGATCAAGATCATCGCCCCG | TTCGTCGTCCTGAGAAGTCG |
| *MYOG* (NM_002479.5) | TCAGCTCCCTCAACCAGGAG | TCTGTAGGGTCAGCCGTGAG |
| *TBP* (NM_003194.4) | TTGTACCGCAGCTGCAAAAT | CGTGGTTCGTGGCTCTCTTA |
| *EEF2* (NM_001961.3) | CCTTGTGGAGATCCAGTGTCC | CTCGTTGACGGGCAGATAGG |

**Supplementary Figure 1**: lower limb muscle MRI in FIN1.

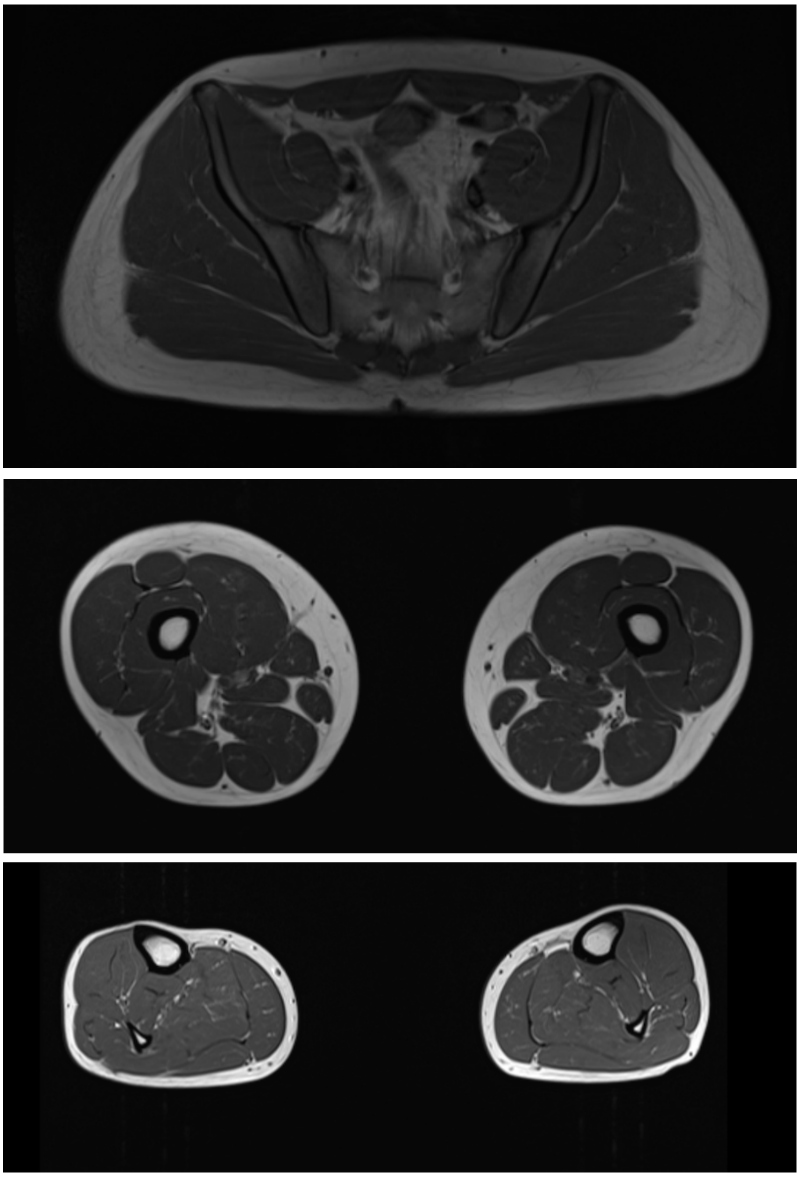

**Supplementary Figure 2**: *OBSCN* LoF variants in gnomAD prioritised by allele frequency. All LOF variants for which there are homozygous individuals have been flagged as “Not LoF” or “Uncertain” in the LoF curation column.

| Variant ID | Source | HGVS Consequence | VEP Annotation | LoF Curation | Clinical Significance | Flags | Allele Count | Allele Number | Allele Frequency | Number of Homozygotes |
|---|---|---|---|---|---|---|---|---|---|---|
| 1-228469903-A-T | E G | p.Arg3252Ter | stop gained | Not LoF | | MNV | 9573 | 275158 | 3.48e-2 | 957 |
| 1-228558992-CCA-C | E G | p.Ser7796Ter | frameshift | Uncertain | | | 740 | 195506 | 3.79e-3 | 4 |
| 1-228559441-GC-G | E G | p.Ser7947ProfsTer82 | frameshift | Uncertain | | | 380 | 226722 | 1.68e-3 | 2 |
| 1-228492263-CA-C | E | p.Arg4755GlyfsTer11 | frameshift | Uncertain | | | 29 | 249174 | 1.16e-4 | 1 |
| 1-228509324-C-T | E | p.Arg5885Ter | stop gained | Uncertain | | | 5 | 248362 | 2.01e-5 | 1 |
| 1-228511027-A-G | E | c.18245-2A>G | splice acceptor | Not LoF | | | 5 | 230190 | 2.17e-5 | 1 |
| 1-228399494-CCA-C | E | p.Gln5ValfsTer31 | frameshift | | | | 1 | 186218 | 5.37e-6 | 0 |
| 1-228399578-C-T | G | p.Gln32Ter | stop gained | | | | 1 | 31310 | 3.19e-5 | 0 |
| 1-228399613-G-A | G | p.Trp43Ter | stop gained | | | | 1 | 31336 | 3.19e-5 | 0 |
| 1-228399729-GC-G | E | p.Arg83AlafsTer6 | frameshift | | | | 1 | 164026 | 6.10e-6 | 0 |
| 1-228399730-CCG-C | E | p.Arg85GlnfsTer67 | frameshift | | | | 2 | 163832 | 1.22e-5 | 0 |
| 1-228399802-CG-C | G | p.Glu107SerfsTer35 | frameshift | | | | 1 | 30988 | 3.23e-5 | 0 |
| 1-228399806-C-T | E | p.Gln108Ter | stop gained | | | | 1 | 130208 | 7.68e-6 | 0 |
| 1-228400030-C-G | E | p.Tyr182Ter | stop gained | | | | 1 | 8182 | 1.22e-4 | 0 |
| 1-228400037-CG-C | G | p.Arg185ProfsTer15 | frameshift | | | | 1 | 25324 | 3.95e-5 | 0 |
| 1-228400196-T-TCACC... | E | p.Gly243AlafsTer12 | frameshift | | | | 1 | 221718 | 4.51e-6 | 0 |
| 1-228400228-GGTGA-G | E | p.Val249LeufsTer12 | frameshift | | | | 1 | 234470 | 4.26e-6 | 0 |
| 1-228400236-AAG-A | E | p.Gly252GlnfsTer103 | frameshift | | | | 1 | 236760 | 4.22e-6 | 0 |
| 1-228400237-A-AG | E | p.Lys253GlnfsTer103 | frameshift | | | | 1 | 236988 | 4.22e-6 | 0 |
| 1-228400241-A-ACC | E | p.Lys253ThrfsTer10 | frameshift | | | | 1 | 237922 | 4.20e-6 | 0 |
